# A Practice on Antibody Hydrophobic Interaction Chromatography Retention Time Prediction using Pre-Trained Large Language Model Fine-Tuning

**DOI:** 10.64898/2026.08.05.742939

**Authors:** Bo Wang, Baowei Cai, Huaxin Chen, Hao Xia, Bo Wang, Jian Liu, Ling Han, Ruwei Wang

## Abstract

Hydrophobicity is a critical property associated with the risk of non-specific binding, and it is commonly assessed using hydrophobic interaction chromatography retention time. Several computational approaches have been developed to predict antibody developability based on pre-trained language models. Such models can be fine-tuned with limited labeled antibody sequences and, in principle, do not require structural information, which is often challenging to obtain. Nevertheless, few studies have achieved strong performance in hydrophobicity prediction without incorporating structural features. Here, we present a case study of fine-tuning the pre-trained model IgBert to predict antibody hydrophobicity. Using Herceptin as a reference, we performed hydrophobic interaction chromatography retention time experiments and generated Herceptin-adjusted datasets. The fine-tuned model achieved a best R^2^ of 0.916, underscoring the critical role of rigorous data quality control. We also synthesized and validated 20 commercially available antibody sequences, and the results showed that the predicted hydrophobic properties were correctly reflected. Our findings provide practical guidance and highlight considerations for future applications of fine-tuned pre-trained language models in antibody hydrophobicity prediction.

**Graphical Abstract:** 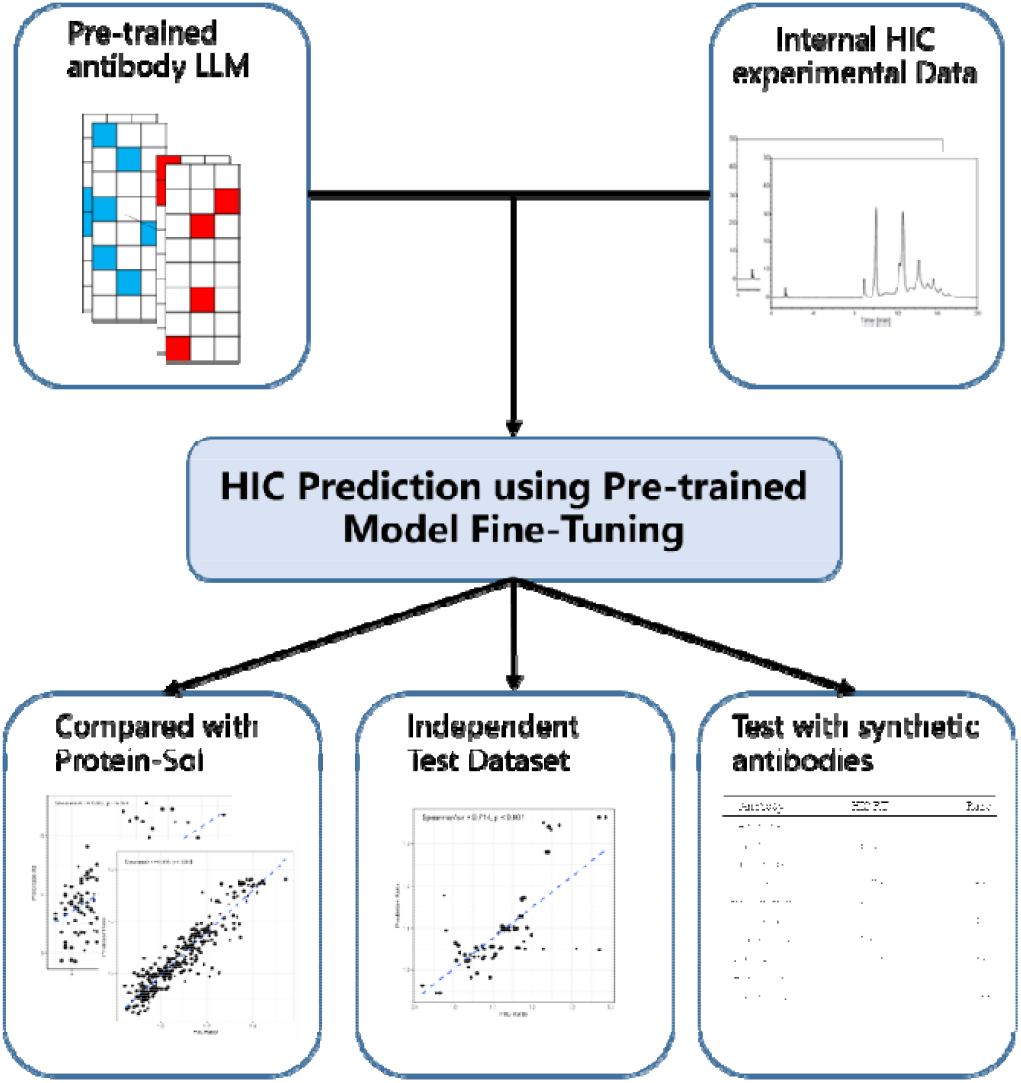

## 1 Introduction

Monoclonal antibodies (mAbs) have become a major therapeutic modality in modern medicine. As of 2023, nearly 200 antibody-based therapeutics have been approved worldwide or are under regulatory review, with over 1,100 candidates currently in Phase I or Phase II clinical trials [1, 2]. Approximately 70% of these investigational antibodies target cancers, highlighting their central role in oncology drug development [1-4]. Despite their success, advancing an antibody candidate from discovery to a marketed product remains a costly and high-risk process, with high attrition rates in late-stage development [5, 6].

To mitigate this risk, early-stage assessment of developability—a collection of molecular properties influencing manufacturability, stability, and clinical performance—is essential [7]. Key developability attributes include binding affinity, thermostability, viscosity, immunogenicity, and hydrophobicity [8]. Among these, hydrophobicity is particularly critical because excessive hydrophobic interactions can lead to non-specific binding and accelerated clearance, thereby reducing serum half-life [6, 9]. Hydrophobicity is routinely quantified by hydrophobic interaction chromatography (HIC), where the retention time (RT) serves as a widely adopted proxy [9].

Computational tools have been developed to facilitate in silico evaluation of HIC RT and related properties. For instance, Protein-Sol predicts solubility directly from sequence features using machine learning [10, 11]; Therapeutic Antibody Profiler (TAP) employs homology modeling and scoring metrics to assess hydrophobicity [12]; and PROPERMAB integrates molecular descriptors with machine learning models to predict HIC RT and viscosity [13]. Other methods explicitly rely on structural descriptors. Hanke et al. introduced the dimensionless retention time (DRT) to estimate HIC RT from structure-derived features [14], while Jain et al. applied random-forest algorithms to predict HIC RT based on side-chain surface exposure in variable regions [9].

More recently, advances in protein language models have enabled data-driven predictions of developability attributes. These models leverage vast unlabeled protein sequence datasets to learn generalizable representations, which can then be fine-tuned with relatively small amounts of labeled antibody data [15]. Several studies have explored their application to developability prediction. AbPROP combines pre-trained sequence embeddings with structural information to predict binding and developability features [16], while AbLEF integrates ensemble language models with structural descriptors to predict HIC RT and aggregation temperature [17]. However, despite these advances, predictive performance for HIC RT remains modest, with reported R^2^ values around 0.5. Importantly, reliance on structural inputs limits scalability, as high-resolution structural data are often unavailable in early-stage antibody discovery [18].

Here, we present a case study of fine-tuning the antibody-specific language model IgBert to predict HIC RT using only sequence information. We experimentally generated HIC RT data for candidate antibodies, adjusting each measurement against Herceptin as an internal control. Fine-tuning on these quality-controlled datasets enabled the model to achieve a best R^2^ of 0.916. These results underscore the importance of rigorous data adjustment and highlight the potential of fine-tuned pre-trained language models to provide accurate, sequence-based predictions of antibody hydrophobicity. We also synthesized and validated 20 commercially available antibody sequences, and the results showed that the predicted hydrophobic properties were correctly reflected. Our findings offer practical guidance for incorporating language models into early-stage antibody developability assessment.

## 2 Methods

### 2.1 Hydrophobic Interaction Chromatography

Analysis was performed on a e2695 Separations Module HPLC system (Waters Corporation). Antibody samples (30µL, 1µg/ml) were injected onto an MAbPac™ HIC-10 HPLC column (4.6 × 100 mm, 5 μm, 100 Å; ThermoFisher Corporation) using a gradient of mobile phase A (1.5 M ammonium sulfate in 50 mM PB, pH 7.0) and B (50 mM PB, pH 7.0) at a flow rate of 1 mL/min. Use Herceptin as a control sample.

### 2.2 Synthetic antibodies

All mAbs were expressed as IgG1 isotype and expressed in CHO cells. The VH and VL encoding gene fragments were subcloned into heavy- and light-chain pcDNA 3.4+ vectors. The plasmid construction and the expression and purification of mAbs were both completed by Genewiz.(Azenta Life Sciences).

### 2.3 Dataset Description

The internal dataset comprised 359 antibody sequences (DataSet-359), each paired with experimentally measured hydrophobic interaction chromatography (HIC) retention time (RT). For every antibody, a matched Herceptin HIC RT measurement was generated in the same experimental batch and served as an internal control. Two training targets were considered: the raw HIC RT values and the Herceptin-adjusted HIC RT values, defined as the ratio of the antibody’s RT to the corresponding Herceptin RT. To confirm the performance of the fine-tuned model, we collected an additional batch of independent test sets consisting 75 antibodies (DataSet-75).

To further validate the model, we incorporated two external public datasets. DataSet-135 consists 137 clinical-stage antibodies that had either received FDA approval or had progressed to Phase II/III clinical trials [8], and consistent with previous reports, two sequences with unusually high HIC RT values (25 min) were identified as suspected outliers and excluded from further analysis, leaving 135 antibodies for training and validation. DataSet-348 consists 348 B cell-derived mAbs with experimental HIC RTs[19].

In addition, the dataset including 20 FDA approval or clinical-stage therapeutic antibodies were synthesized (DataSet-20). These molecules were processed under the same experimental workflow, with HIC RT measured alongside Herceptin controls, thereby ensuring comparability to the internal dataset. The antibody sequences, and the experimental and predicted results are in the **Supplementry File**.

### 2.4 Pre-Trained Language Model and Model Fine Tuning

We adopted IgBert, a pre-trained antibody language model trained on more than two billion unpaired antibody sequences and approximately two million paired light–heavy chain sequences from the Observed Antibody Space (OAS) database [18-20]. The model consists of 30 transformer layers, each with a hidden dimension of 1024, totaling approximately 420 million parameters. Previous benchmarking has demonstrated that IgBert outperforms existing antibody- and protein-focused language models across multiple fine-tuning tasks related to antibody developability. However, the task of hydrophobicity prediction was not evaluated and practiced in the original study [18].

For hydrophobic interaction chromatography (HIC) retention time (RT) prediction, a regression head was added on top of the pre-trained backbone. The regression output was derived from a fully connected layer appended to the pooled representation. Model fine-tuning was performed using the internal training dataset. All scripts were implemented in Python 3.8.20, with dependencies including PyTorch, Transformers, scikit-learn, NumPy, and pandas.

Training was conducted for up to 40 epochs with a batch size of 32 and an initial learning rate of 1e-5. A learning rate scheduler decreased the rate by a factor of 0.2 every 10 epochs. Early stopping was employed to prevent overfitting, with a patience of three epochs and a minimum loss improvement (delta) of 0.01. During each epoch, the dataset was randomly split into 90% for training and 10% for validation, effectively implementing a repeated 10-fold cross-validation strategy.

Model performance was evaluated using several statistical metrics, including mean squared error (MSE), root mean squared error (RMSE), mean absolute error (MAE), coefficient of determination (R2), and training loss. The codes are accessible at https://github.com/AlkaidWang/IgBert2HIC.

### 2.5 Performance Comparisons

Both raw HIC RT values and Herceptin-adjusted HIC RT ratios were used as training targets during fine-tuning, and the corresponding validation metrics were compared to assess the impact of internal control adjustment.

For benchmarking, the widely used sequence-based tool Protein-Sol [10, 11] was deployed locally. Protein-Sol is comparable to our approach because it only requires sequence information without structural features. The tool was evaluated on both the internal dataset and the external clinical dataset. However, direct comparison between the fine-tuned model and Protein-Sol on the public dataset was not feasible, as Protein-Sol predicts raw HIC RT values whereas our fine-tuned model outputs Herceptin-adjusted ratios.

Finally, the fine-tuned model was applied to the synthetic antibody dataset (20 molecules). Predictions were compared with Herceptin-adjusted experimental HIC RT values. Performance was quantified using the Spearman correlation coefficient to capture the rank-ordering agreement between predicted and observed values.

## 3 Results

### 3.1 Datasets Comparison

We first examined the distributions of HIC RT values across the datasets, and the descriptive statistics are summarized in **Table 1**. For Herceptin, the measured HIC RT ranged from 11.87 to 18.45 minutes with a standard deviation of 1.27 minutes, reflecting systematic variability attributable to experimental factors such as reagents, instrumentation, and operator differences. This observation suggests that model training based directly on raw HIC RT values may be affected by batch-specific noise.

**Table 1.** The range, average, median, standard deviation values of datasets.

| Dataset | Type | Min. | Max. | Mean | Median | SD. |
| --- | --- | --- | --- | --- | --- | --- |
| DataSet-359 | Herceptin (min.) | 11.87 | 18.45 | 15.61 | 15.84 | 1.27 |
|  | Antibodies (min.) | 10.63 | 25.01 | 16.82 | 16.72 | 2.30 |
|  | Adjusted Ratio | 0.84 | 1.55 | 1.08 | 1.06 | 0.14 |
| DataSet-20 | Antibodies (min.) | 14.22 | 25.28 | 20.15 | 20.24 | 3.54 |
|  | Adjusted Ratio | 0.90 | 1.60 | 1.27 | 1.28 | 0.22 |
| DataSet-135 | Antibodies (min.) | 8.50 | 13.70 | 10.11 | 9.80 | 1.10 |
| DataSet-348 |  | 8.50 | 13.90 | 9.39 | 9.10 | 0.83 |

The internal dataset displayed a markedly broader distribution of antibody HIC RTs (10.63–25.01 min.) while the public dataset ranged from 8.50 minutes to 13.90 minutes. This discrepancy implies that the internal and public datasets may have been generated under different experimental systems or protocols. Consequently, machine learning models trained exclusively on public data—for example, Protein-Sol—may exhibit limited generalizability when applied to our internal measurements.

In addition, the public datasets lack an internal reference standard such as Herceptin, precluding the possibility of ratio-based adjustment. To address this, several antibodies from the public dataset were synthesized and experimentally tested alongside Herceptin controls. These measurements enabled calculation of Herceptin-adjusted HIC RT ratios, which in turn allowed consistent evaluation of both our fine-tuned model and sequence-based public tools such as Protein-Sol.

### 3.2 Fine-tuned Model Training

The fine-tuned antibody model reached convergence after 8 epochs. Both Loss and MSE decreased to values close to 0, while RMSE and MAE were reduced to below 0.1. Notably, the coefficient of determination (R^2^) obtained from 10-fold cross-validation was 0.63, which is higher than the best performance reported for AbLEF (R^2^ = 0.49 [17]). The detailed validation parameters are shown in **Supplementary File**. In contrast, when the model was fine-tuned using the raw HIC retention times from the internal dataset under the same training parameters, the performance was substantially worse, with Loss of 8.55, MSE of 6.60, RMSE of 2.57, MAE of 1.99, and a negative R^2^ of -0.18. These results suggest that systematic errors in the raw experimental data can significantly impair the effectiveness of model fine-tuning.

### 3.3 Model Performance Evaluation

We evaluated Protein-Sol using both the public and internal antibody datasets by comparing its predicted HIC RTs with the experimental values. The correlation coefficient was 0.66 **(Fig 1A)** for the DataSet-135 and 0.41 **(Fig 1B)** for the DataSet-348, while with the internal DataSet-359, the coefficent was only 0.17 for RTs and 0.20 for ratios **(Fig 1C, 1D)**. The relatively strong correlation with the public dataset can be attributed to the fact that Protein-Sol was trained on the same set of 137 clinical antibodies. However, the prediction completely failed on the internal dataset, likely due to the limited size of the original training set, which reduces the model’s generalizability to unseen sequences. In contrast, the fine-tuned antibody model achieved a correlation coefficient of 0.916 on the internal dataset **(Fig 2)**. Using the independent test set (DataSet-75), the correlation coefficient dropped to 0.714, which is still higher than that of Protein-Sol **(Fig 3)**. Furthermore, in actual production, the predicted values are converted into qualitative indicators, which can be used for inferior-exclusion screening. It can be seen from **Fig 3** that the hydrophobicity between 0.9 and 1.3 is more appropriate. Although the predicted values outside the range have outliers, they do not affect the qualitative judgment.

**Fig. 1.**
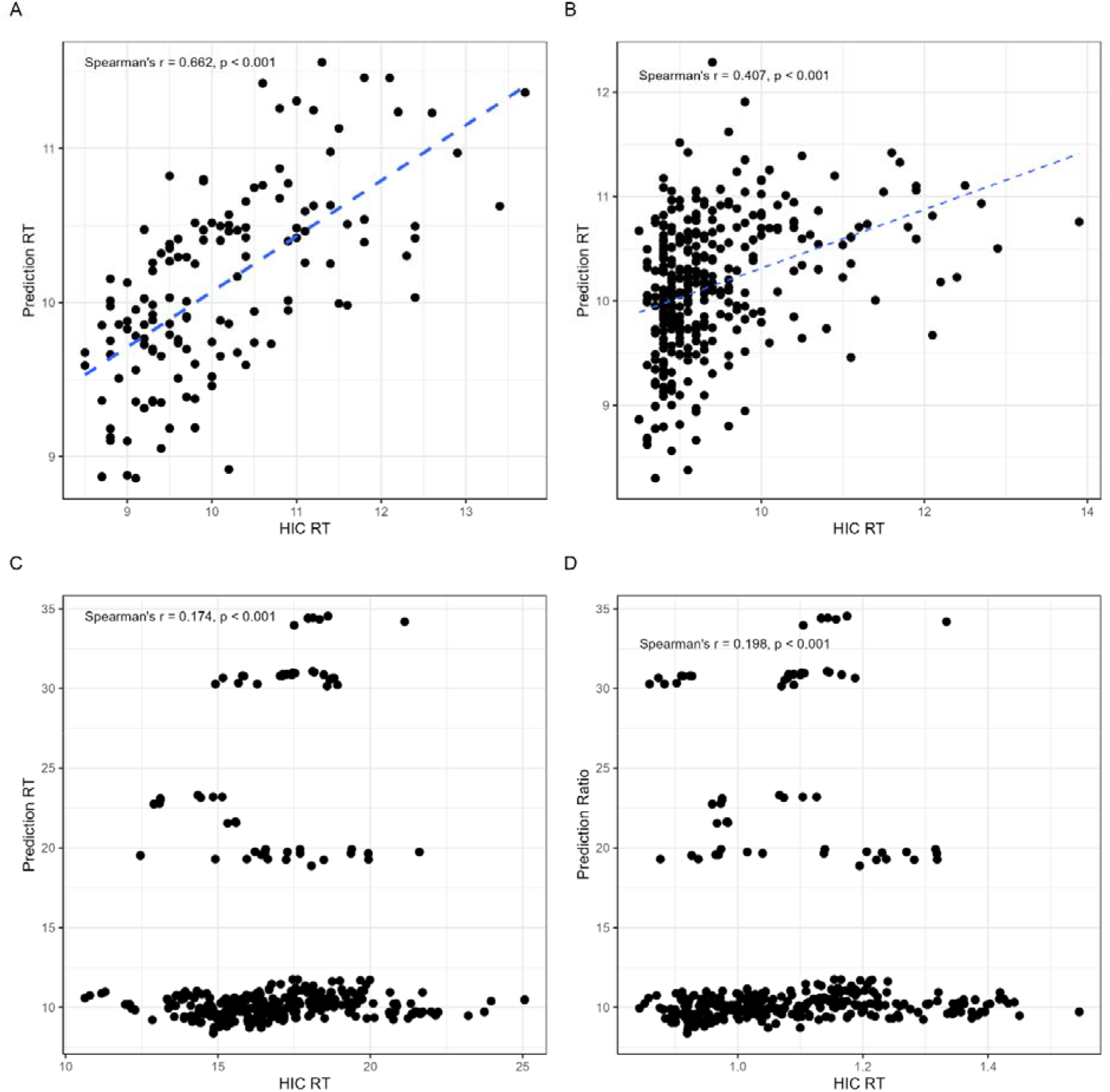
Performance evaluation of Protein-Sol. (**A**) Using the DataSet-135, which are the training set of Protein-Sol, this model got a correlation coefficient of 0.662, between experimental HIC RTs and the predicted ones. (**B**) However, using the public DataSet-348, the correlation coefficient dropped to 0.407. (**C, D**) Using the internal DataSet-359, Protein-Sol failed to predict the RTs accurately, with the correlation coefficient of 0.174 and 0.198 for RTs and Ratios correspondingly.

**Fig. 2.**
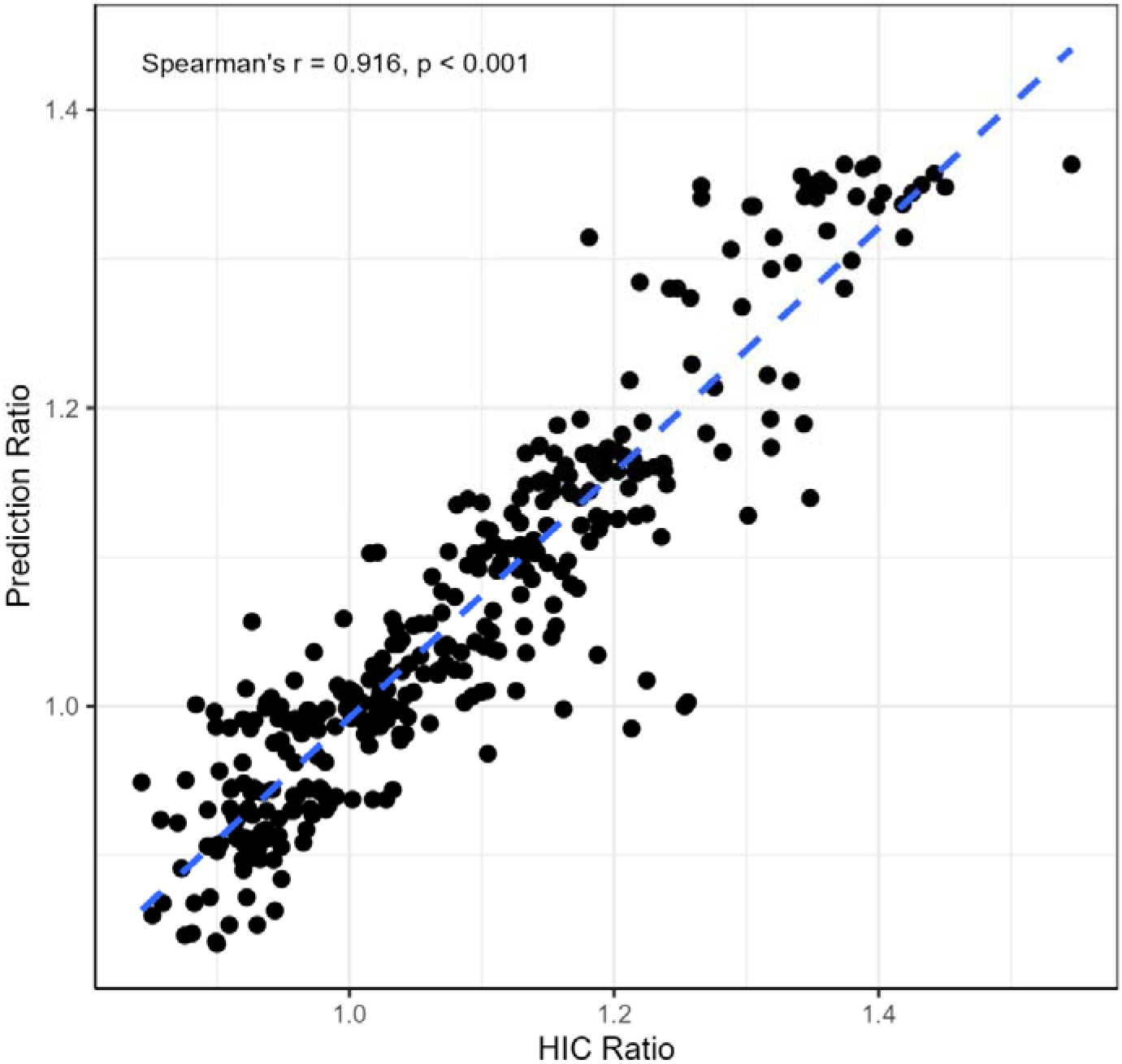
Using DataSet-359 to evaluate the performance of fine-tuned model, which is the training set of this model, it got the best correlation coefficient of 0.916.

**Fig. 3.**
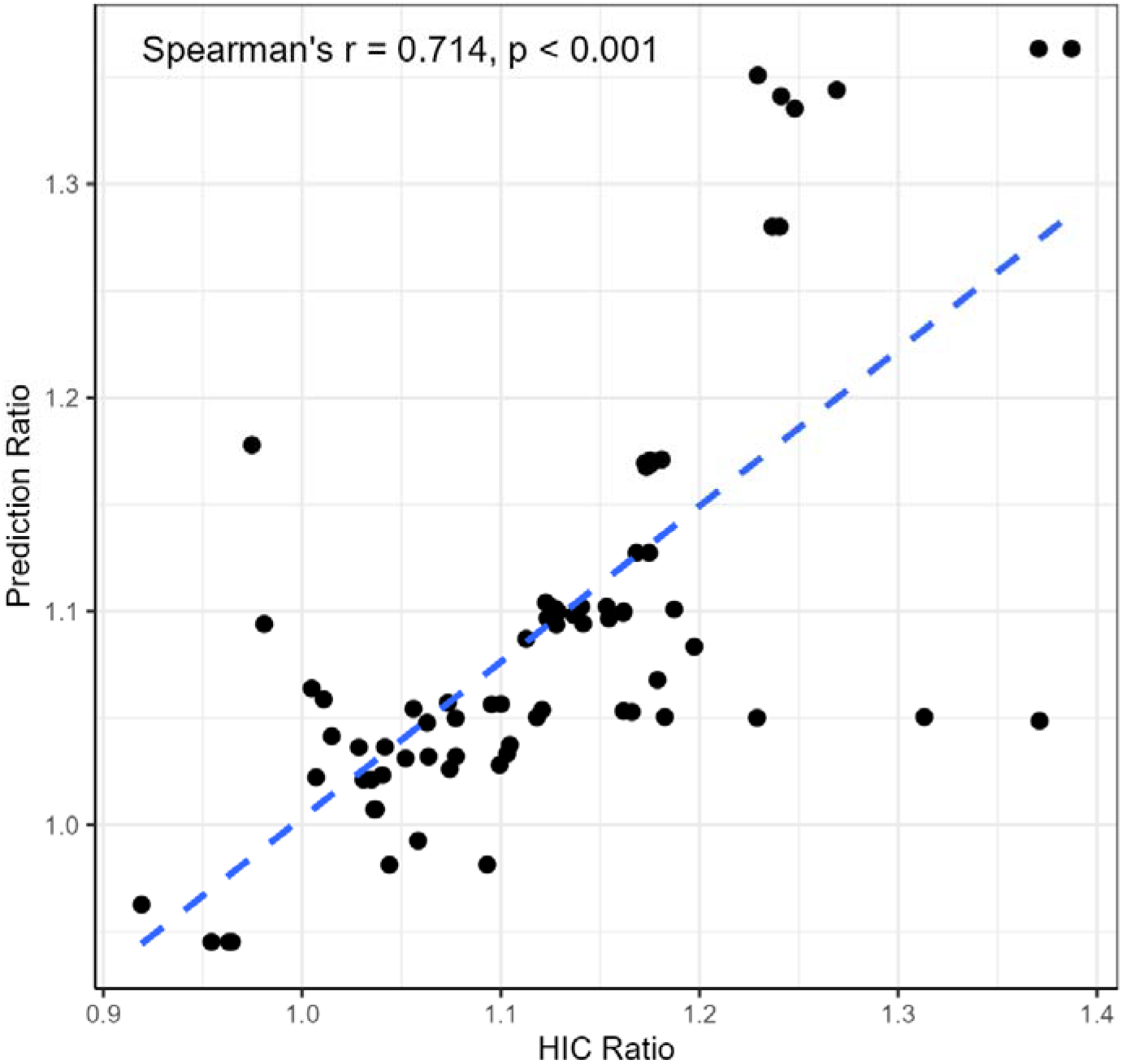
Using DataSet-75 to confirm the performance of fine-tuned model, which is the independent test set, the correlation coefficient dropped to 0.714, but the outliers were mostly the bad ones (out of the range of 1.0-1.2).

### 3.4 Synthetic Data Set Validation

To confirm whether the performance of the fine-tuned model was dependent on the training dataset, we performed tests with clinical-stage antibodies. Twenty antibodies were synthesized and tested for HIC RT with the Herceptin as internal control internally, and the detailed results are shown in **Table 2**. The predicted ratios ranges from 0.976 to 1.246, which suggests that this range can be used in pratical screening.

**Table 2.** The experimental and predicted RT/Ratio values of the 20 synthesized clinical-stage antibodies.

| Antibody | HIC RT | Ratio | Predicted Ratio |
| --- | --- | --- | --- |
| veltuzumab | 20.237 | 1.277 | 1.246 |
| daclizumab | 16.177 | 1.021 | 1.222 |
| rituximab | 19.83 | 1.251 | 1.215 |
| visilizumab | 14.218 | 0.897 | 1.209 |
| tralokinumab | 16.262 | 1.026 | 1.164 |
| galiximab | 22.565 | 1.424 | 1.145 |
| mavrilimumab | 23.892 | 1.507 | 1.126 |
| ponezumab | 21.258 | 1.341 | 1.125 |
| evolocumab | 17.303 | 1.092 | 1.111 |
| rilotumumab | 22.794 | 1.438 | 1.104 |
| lirilumab | 25.283 | 1.595 | 1.100 |
| seribantumab | 19.812 | 1.250 | 1.096 |
| atezolizumab | 24.778 | 1.563 | 1.089 |
| ranibizumab | 22.792 | 1.438 | 1.063 |
| tanezumab | 20.045 | 1.265 | 1.051 |
| sirukumab | 20.247 | 1.277 | 1.045 |
| gevokizumab | 14.678 | 0.926 | 1.043 |
| pembrolizumab | 24.015 | 1.515 | 1.024 |
| codrituzumab | 14.507 | 0.915 | 1.024 |
| lebrikizumab | 22.297 | 1.407 | 0.976 |

## 4 Discussion

We developed and evaluated a sequence-only strategy for predicting antibody hydrophobicity by fine-tuning a pre-trained antibody language model (IgBert [18]) on experimentally measured HIC retention times (RT) that were adjusted by the internal control (Herceptin). Our principal finding is that internal-control adjustment substantially improves model convergence and predictive accuracy: models trained on Herceptin-adjusted RT ratios markedly outperformed models trained on raw RT values, which were strongly affected by batch-specific variability. This observation highlights that reducing systematic assay variation is essential when using real-world biophysical measurements for model fine-tuning.

The impact of assay variability and normalization on downstream statistical modeling is well documented in omics and high-throughput biophysics: appropriate normalization or use of internal standards reduces unwanted technical variation and improves comparability across batches and platforms [21]. In our experiments, Herceptin served as a consistent internal control in each run and enabled calculation of adjusted RT ratios that were much less sensitive to instrument- or operator-dependent shifts than raw RT values.

When benchmarked against an established sequence-only tool (Protein-Sol [10, 11]), our fine-tuned model showed superior performance on the internal dataset while Protein-Sol only correlated well with the public dataset on which it was developed. This contrast underscores two points: (i) tools trained on relatively small or homogeneous public datasets may generalize poorly to measurements collected under different experimental conditions, and (ii) combining broad pre-training (large sequence corpora) with careful, internally adjusted fine-tuning can produce models that generalize to in-house experimental systems.

Our use of a clinical-stage antibody reference dataset and synthesis/measurement of selected public antibodies further supports the model’s practical utility. The public clinical dataset we used (137 clinical-stage antibodies) is a commonly used benchmark in antibody developability studies and captures typical biophysical ranges of late-stage antibodies [8]; however, those public measurements do not include internal controls, which limits direct cross-study normalization.

The effectiveness of sequence-based pre-training followed by task-specific fine-tuning is consistent with recent work (FLAb2) showing that large protein language models learn representations that implicitly encode structural and functional information from sequence alone, enabling strong zero- and few-shot generalization across diverse protein tasks[20]. However, results for hydrophobisty with few-shot models were omitted due to a lack of data. In our setting, the combination of (i) an antibody-focused pre-trained model, (ii) carefully curated and internally adjusted HIC RT labels, and (iii) appropriate fine-tuning hyperparameters produced a model with high predictive fidelity (internal R^2^ and strong rank correlation). It shows the importance of training dataset.

Nevertheless, important limitations and practical considerations remain. Although fine-tuning requires far fewer labeled examples than training from scratch, the labeled data that are used must be high quality and consistent: pooling measurements across different experimental platforms without appropriate normalization can degrade model performance. This challenge—data heterogeneity and limited labeled sets—has been recognized more generally in AI applications to biology and medicine and remains a central obstacle to robust model deployment [22]. Besides, although our internal control approach reduced batch effects and improved model performance, scaling to industrial pipelines will require systematic procedures for (i) routine inclusion of stable internal controls, (ii) metadata capture (buffer composition, column lot, instrument settings), and (iii) shared normalization standards so models trained in one lab can be adapted or transferred to another [23].

Looking forward, sequence-only fine-tuning of large protein language models can be deployed in multiple stages of antibody discovery and development. Immediate applications include triage of large candidate libraries to prioritize molecules with favorable hydrophobicity profiles, incorporation into multi-parameter developability scoring pipelines, and early-stage optimization for formats such as antibody-drug conjugates where hydrophobicity critically affects conjugate stability and pharmacokinetics. The method may also accelerate iterative “design–make–test” cycles when coupled with high-throughput experimental assays, reducing the time and cost of lead optimization. Advances in foundation models for proteins and multi-task learning suggest a future in which generalized models can support several developability endpoints simultaneously, while transfer learning allows lab-specific calibration with modest labeled datasets [15].

## 5 Conclusions

In conclusion, our work demonstrates that fine-tuning an antibody-specific language model on internally normalized HIC RT measurements yields accurate sequence-only predictions of hydrophobicity. The main practical lesson is that careful experimental design—particularly inclusion of an internal control and rigorous normalization/adjustment — can be as important as model architecture for achieving reliable predictive performance. We anticipate that combining high-quality experimental datasets with pre-trained sequence models will become an increasingly practical route to integrate AI into developability workflows and to reduce late-stage attrition in antibody development.

## Supplementary Information

The online version containsSupplementary material available at https://.

## Funding

All authors are/were Yangtze River Pharmaceutical Group employees when conducting the work. The work here was fully funded by Yangtze River Pharmaceutical Group.

## Data availability

The raw data used for fine-tuned training during the current study is only available from the corresponding author on reasonable request. The fine-tuned model weight file is avaialbe at https://github.com/AlkaidWang/IgBert2HIC.

## Table and Figure Captions

**Table 1** The distributions of HIC RT/Ratio values across the datasets.

**Table 2** The experimental and predicted RT/Ratio values of the 20 synthesized clinical-stage antibodies.

## Notes

https://github.com/AlkaidWang/IgBert2HIC

### Competing Interest Statement

The authors have declared no competing interest.

